# XDec Simplex Map of Breast Cancer Cell States Enables Precise Modeling and Targeting of Breast Cancer

**DOI:** 10.1101/2022.07.06.498858

**Authors:** Oscar D. Murillo, Varduhi Petrosyan, Emily L. LaPlante, Lacey E. Dobrolecki, Michael T. Lewis, Aleksandar Milosavljevic

## Abstract

The characterization of cancer cell states within the tumor microenvironment is a key to understanding tumor biology and an important step toward the development of precision therapies. To reconstruct this information from bulk RNA-seq profiles, we developed the XDec Simplex Mapping (XDec-SM) approach, a reference-optional deconvolution method that leverages single-cell information, when such information is available, to map tumors and the states of constituent cells onto a biologically interpretable, low-dimensional space. When applied to breast tumors in The Cancer Genome Atlas (TCGA), XDec-SM infers the identity of constituent cell types and their proportions. XDec-SM also infers cancer cells states within individual tumors that associate with DNA methylation patterns, driver somatic mutations, pathway activation and metabolic coupling between stromal and breast cancer cells. By projecting tumors, cancer cell lines, and PDX models onto the same map, we identify both *in vitro* and *in vivo* models with matching cancer cell states. Map position is also predictive of therapy response, thus opening the prospects for precision therapy informed by experiments in model systems matched to tumors *in vivo* by cancer cell state.

## INTRODUCTION

Molecular profiling of breast tumors over the past two decades has further reinforced the understanding of breast cancer as a highly heterogeneous disease with a staggering complexity of molecular aberrations at both the tissue and cellular levels. Large data banks such as The Cancer Genome Atlas (TCGA) (Liu et al., 2018) and the International Cancer Genome Consortium (ICGC) (Campbell et al., 2020) characterize a vast array of tumors. However, the multi-omics characterization of these tumors is confounded by the bulk profiling analyses. Despite the advances in sequencing that allow for the high throughput processing of these highly complex samples, the variability in tumor composition averages cell type specific signatures (Beca and Polyak, 2016; Place et al., 2011). The averaging also precludes access to the state of cancer cells within tumors, which is essential for understanding effects of somatic mutations, identifying interactions between cancer cells and the tumor microenvironment and for predicting therapy response.

Single-cell profiling provides direct access to the states of constituent cell types within individual tumors. Among the single-cell methods, the most widely used is single-cell RNA sequencing (scRNA-seq), (González-Silva et al., 2020; Karaayvaz et al., 2018; Patel et al., 2014; Puram et al., 2017). While in principle providing the ultimate level of resolution, the scRNA-seq method has limitations: the depth of coverage is relatively sparse precluding precise quantitation; biased towards highly abundant cell types; the cost per sample is high (Chen et al., 2019; Haque et al., 2017; Ilicic et al., 2016; Kolodziejczyk et al., 2015); and it is not readily applicable to the formalin-fixed paraffin-embedded (FFPE) samples which are routinely collected in practice. Computational deconvolution is highly synergistic with single-cell profiling, as it decreases cost and technical requirements dramatically, while benefiting from the information about the diversity of cell types gathered by scRNA-seq. Methods that combine deconvolution and single-cell profiling such as CIBERSORTx (Newman et al., 2019) and MuSiC (Wang et al., 2019) have demonstrated the synergy of the two approaches. However, these computational deconvolution methods are reference-based in that they explain bulk profiles as linear combinations of *a priori* defined profiles of constituent cell types that have previously been physically isolated, thus precluding data-driven discovery of recurrent states of cancer cells and their comparison to established tumor classification (e.g., Luminal, Basal, and HER2 breast cancer subtypes). Although methods such as EcoTyper (Luca et al., 2021) can identify cell states, these methods still rely on cell type references. One key limitation of the reference-based approach is that the physical separation that is needed for obtaining the references perturbs the constituent cell states, thus calling for deconvolution strategies that can infer cell states (beyond those observable in single-cell references) from bulk profiling data.

Despite the advances in both epigenomic and transcriptomic computational deconvolution, no deconvolution method has been shown to estimate cell states that correspond to subtypes of breast cancer or other solid tumors in the context of the complete tumor microenvironment. To address this knowledge gap, we developed XDec-SM a new methodological framework and algorithm based on simplex mapping that combines bulk RNA-seq profiles of individual tumors and publicly accessible relevant scRNA-seq information in a way that enables discovery of new recurrent states of cancer cells. The method is reference-optional as it may leverage information from references when available (e.g., scRNA-seq profiles). If references are provided. Moreover, the method only requires them the purpose of identifying informative loci (e.g., genes, methylation loci). for deconvolution. Additionally, information about informative loci from an external source other than the references may also be used in conjunction (e.g., literature-based signature).

We apply XDec-SM to breast tumors in the TCGA collection to construct the first simplex map of breast cancer, defined by the cellular makeup of individual tumor samples and the state of constituent cell fractions. Because the algorithm is reference-optional, it produces data-driven estimates of constituent cell types using both single cell information and previously published PAM50 subtype loci (Campbell et al., 2020; Nielsen et al., 2010; Parker et al., 2009). We determine associations between the cancer cell state, DNA methylation patterns, somatic driver mutations, activation of cancer promoting pathways and heterotypic interactions within the tumor microenvironment. The map enables precise modeling of tumors by placing the deconvoluted *in vivo* cancer cells, cancer cell lines, and PDX models within the same low-dimensional coordinate system and enables the use of experimentally tractable models to predict tumor-specific response to therapy. To empower future studies, we deploy XDec-SM as a R package. We also make it accessible on-line via a web user interface for placing individual breast tumors, PDX, and cell line models on the map based on their bulk RNA-seq profiles.

## RESULTS

### XDec-SM deconvolution allows for the mapping of BRCA tumors onto a cancer cell state map

The XDec-SM deconvolution method (STAR Methods) was applied to deconvolute the bulk RNA-seq breast cancer tumor profiles from the TCGA collection by leveraging publicly available single cell sequencing data (Karaayvaz et al., 2018) (Figure 1A). The informative genes (n = 323) were defined as the union of genes differentially expressed in the scRNA-seq data in different breast tumor cell types and the PAM50 genes (Figure S1A). The deconvolution algorithm identified nine distinct cell types within the tumors (Figure S1B, Figure S1C) explaining 85.2% of the variance. Five distinct epithelial cell types, a Cancer Associated Fibroblast (CAF) as well as CAF adipocyte, macrophage, and a T-cell cell types were identified.

**Figure 1.**
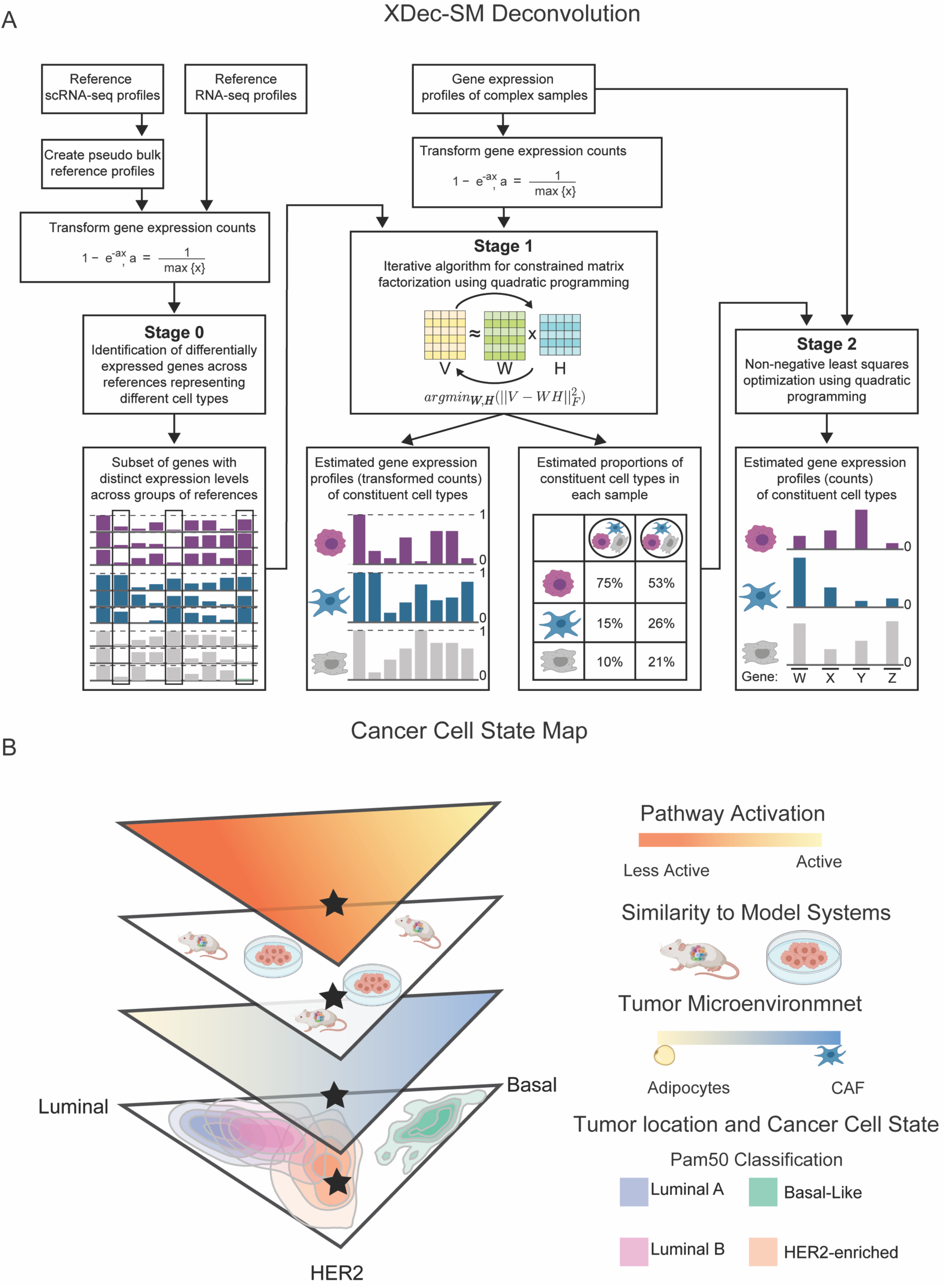
Cancer cell state map built with XDec-SM deconvolution. (A) The XDec-SM algorithm contains three stages (Stage 0, 1, 2). Stage 0 is generalized to utilize a set of reference RNA-seq or scRNA-seq profiles. Reference scRNA-seq profiles with user provided cell-type classifications are utilized to create pseudo bulk reference profiles by summing every five profiles ordered by total gene coverage. The reference RNA-seq profiles or pseudo bulk profiles are transformed to a 0-1 range to identify differentially expressed genes representing different cell types. The subset of informative genes with distinct expression levels across the groups of references are utilized in Stage 1. In Stage 1, using the gene expression profiles of complex samples transformed to 0-1, XDec-SM estimates the gene expression profiles (transformed counts) of constituent cell types and the proportions of constituent cell types in each sample. XDec-SM uses an iterative algorithm for constrained matrix factorization using quadratic programming. The estimated proportions of constituent cell types in each sample are used in Stage 2. In Stage 2, using the untransformed gene expression profiles of the same complex samples and the proportions estimated in Stage 1, XDec-SM estimates the gene expression profiles (counts) of constituent cell types. Stage 2 uses non-negative least squares optimization using quadratic programming. (B) Cancer cell state map of breast cancer. The cancer cell state map of breast cancer acts as a framework to determine a tumor’s cancer cell state (layer 1 bottom). The map is also indicative of a tumor’s microenvironment (layer 2), similarity to model organisms including murine models and cell lines (layer 3), and the activation of cancer associated pathways (layer 4 top).

Unlike other reference-based methods, XDec-SM utilized a reference-optional, data driven approach to define constituent cell types within tumors and is not constrained by the availability of references profiles. In an improvement over the EDec (Onuchic et al., 2016) algorithm, both a normal adipocyte stromal profile and a CAF stromal profile were identified (Figure S1D). Of the five epithelial profiles identified by XDec-SM, three were identified as Luminal, Basal, and HER2 cancer epithelial profiles by observing their high abundance in TCGA tumors with respective Luminal, Basal, and HER2 subtype classification (Figure S1D, STAR Methods). To further validate these designations of deconvoluted profiles, Stage 2 XDec-SM deconvolution was performed, and the cell type specific expression of marker genes for the three subtypes was confirmed (Figure S2, STAR Methods).

While the majority of cancer cells within a specific TCGA tumor was typically of the same type (e.g., Luminal) as its PAM50 classification based on its bulk profile, other subtypes (e.g., Basal, HER2) were also detected albeit in smaller proportions. As we discuss in detail below, this is only partially due to the presence of cancer cells of different type; the fact that the cancer cells can be modeled as a linear combination of pure subtypes also reflects intermediate cancer cell states that resemble to various degrees (as indicated by relative proportion estimates) the three subtypes. In other words, the “relative proportions” of the three subtypes also reflects relative similarities to the “pure” deconvoluted cancer cell states of the three subtypes. Mindful that the deconvoluted proportions of the three subtypes may to a significant degree reflect relative similarity, we placed the cancer cell fractions of each TCGA tumor within a two-dimensional simplex map (Figure 1B). This three-dimensional simplex map acts as a framework onto which breast cancer tumors can be projected, and the position of a tumor in the cancer cell state is associated with the tumor microenvironment, similarity to model organism, and other tumor biology such as the activation of tumor promoting pathways (Figure 1B).

### XDec-SM compares favorably to other deconvolution algorithms

After deconvoluting TCGA breast cancer samples, we next compared XDec-SM against other widely used computational deconvolution methods (STAR Methods). Because CIBERSORTx (Newman et al., 2019) is also equipped to employ scRNA-seq reference profiles and outputs both cell type specific profiles and per-sample proportions, we compared cell type proportions estimated by the two methods (Figure 2A). Despite relatively high correlations (epithelial R^2^ = 0.93; immune R^2^ = 0.80; stromal R^2^ = 0.78, Figure 2A), systematic discrepancies between the two methods could be observed (Figure 2A).

**Figure 2.**
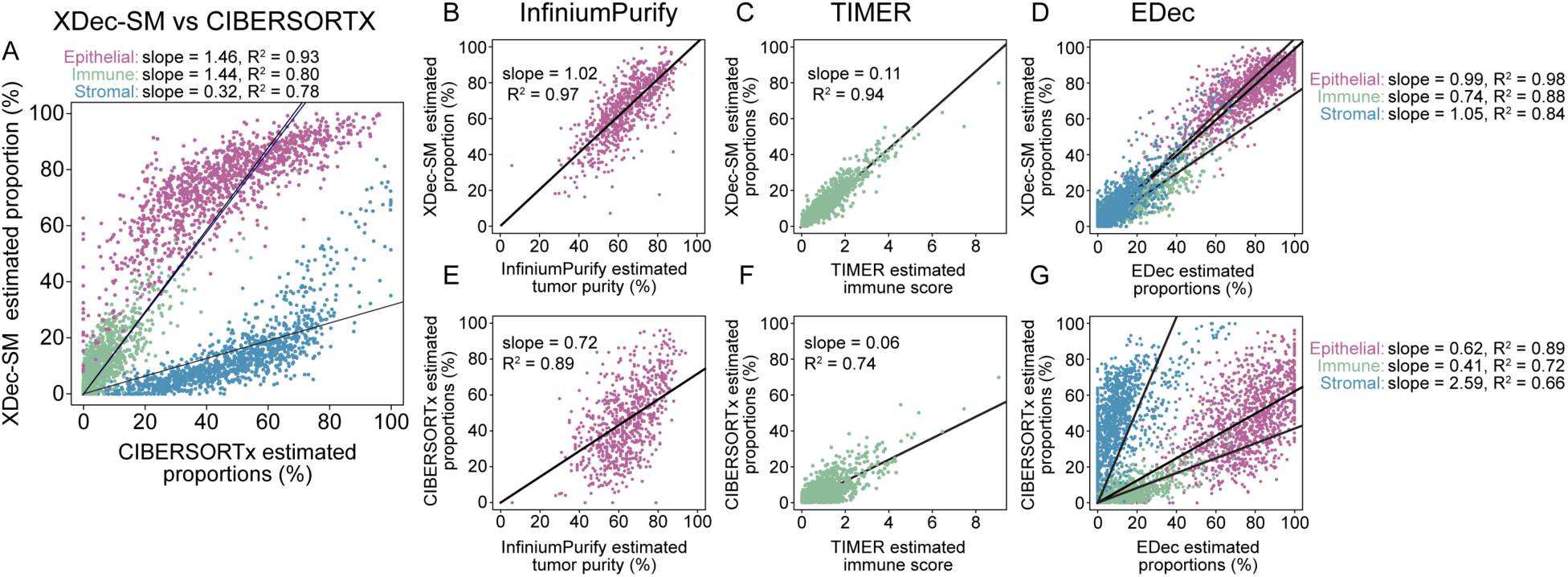
Comparison of XDec-SM estimated proportions to other deconvolution methods. (A) Scatterplot between CIBERSORTx (x-axis) and XDec-SM (y-axis) estimated per-sample proportions of each constituent cell type. For XDec-SM, the five epithelial profiles are summed to represent epithelial, the two stromal profiles are summed to represent stromal, and the T cell and Macrophage are summed to represent immune. For CIBERSORTx, the three immune (B cell, T cell, macrophage) profiles are summed to represent immune and the two stromal (stroma, endothelial) profiles are summed to represent stromal. (B) Scatterplot between InfiniumPurify (x-axis) estimated tumor purity and XDec-SM (y-axis) estimated per-sample proportion of epithelial. For XDec-SM, the five epithelial profiles are summed to represent epithelial. (C) Scatterplot between TIMER (x-axis) estimated immune score and XDec-SM (y-axis) estimated per-sample proportion of epithelial. For XDec-SM, the two immune profiles are summed to represent immune and compared to the sum of all immune subtype scores estimated by TIMER. (D) Scatterplot between EDec (x-axis) and XDec-SM (y-axis) estimated per-sample proportions of each constituent cell type. For EDec, the six epithelial profiles are summed to represent epithelial. For XDec-SM, the five epithelial profiles are summed to represent epithelial, the two stromal profiles are summed to represent stromal, and the T cell and Macrophage are summed to represent immune. (E) Scatterplot between InfiniumPurify (x-axis) estimated tumor purity and CIBERSORTx (y-axis) estimated per-sample proportion of epithelial. (F) Scatterplot between TIMER (x-axis) estimated immune score and CIBERSORTx (y-axis) estimated per-sample proportion of epithelial. For CIBERSORTx, the three immune profiles are summed to represent immune and compared to the sum of all immune subtype scores estimated by TIMER. (G) Scatterplot between EDec (x-axis) and CIBERSORTx (y-axis) estimated per-sample proportions of each constituent cell type. For EDec, the six epithelial profiles are summed to represent epithelial. For CIBERSORTx, the two immune (B cell, T cell, macrophage) profiles are summed to represent immune and the two stromal (stroma, endothelial) profiles are summed to represent stromal.

To interpret the discrepancies between XDec-SM and CIBERSORTx, we proceeded to independently estimate cell type proportions by InfiniumPurify (Zhang et al., 2015) and TIMER (Li et al., 2017). We observed higher concordance between cancer cell fraction estimates of InfiniumPurify and XDec (slope = 1.02, R^2^ = 0.97, Figure 2B), than CIBERSORTx (slope = 0.72, R^2^ = 0.89, Figure 2E). We also observed higher concordance between immune fraction estimates of TIMER and XDec-SM (slope = 0.11, R^2^ = 0.94, Figure 2C) than CIBERSORTx (slope = 0.06, R^2^ = 0.74, Figure 2F). Taken together, higher concordances of XDec-SM with both InfiniumPurify and TIMER suggest that constituent cell proportions predicted by XDec-SM are accurate.

Because XDec-SM was developed using the EDec method as the starting point and focuses on the state of gene activation (vs. gene-specific transcription levels), we next asked if XDec-SM recapitulates the results of epigenomic methylation-based deconvolution. Despite XDec-SM and EDec exploiting different data types – RNA-seq and DNA methylation respectively both methods focus on the gene activation state (constrained to the 0-1 scale). Therefore, we hypothesize that these methods will be concordant. Indeed, we observed high concordance between XDec-SM and EDec across all three cell class proportions (epithelial, slope = 0.99, R^2^ = 0.98; immune, slope = 0.74, R^2^ = 0.88; stromal, slope = 1.05, R^2^ = 0.84, Figure 2D). CIBERSORTx shows lower concordance with EDec (epithelial, slope = 0.62, R^2^ = 0.89; immune, slope = 0.41, R^2^ = 0.72; stromal, slope = 2.59, R^2^ = 0.66, Figure 2G).

### Cancer cell state mapping provides insights into intra-tumoral heterogeneity

Unlike the categorical PAM50 classification of BRCA tumors, the XDec-SM mapping approach places cancer cells on the spectrum between the pure Luminal, Basal and HER2 subtypes (Figure 3A, Figure 3B). As expected, the tumors classified as Basal by PAM50 were clustered primarily in the Basal corner of the simplex map. However, we observed that the tumors classified as Luminal or HER2 by PAM50 were found on a spectrum between the Luminal and HER2 vertices.

**Figure 3.**
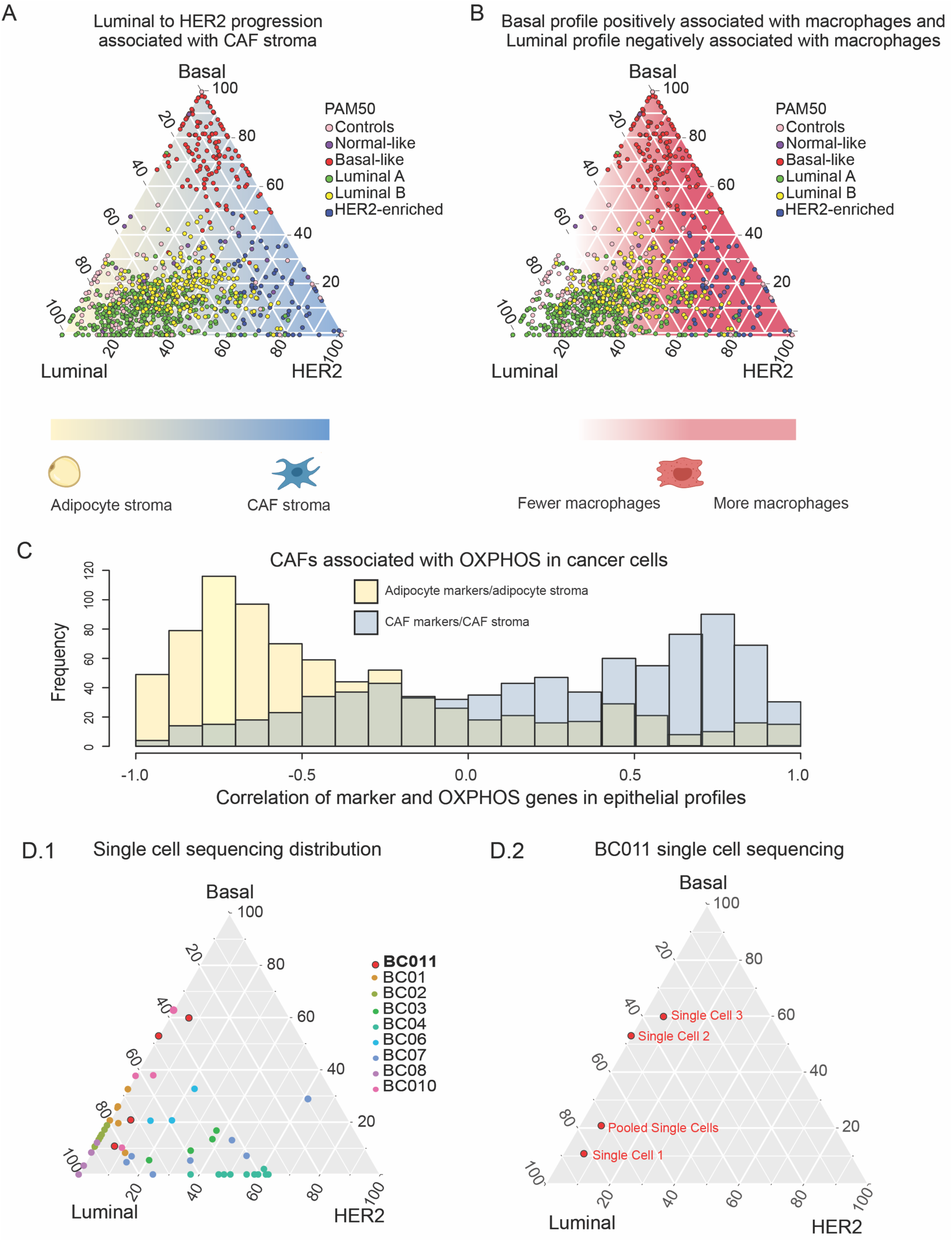
Cancer cell state map reveals intra-tumoral heterogeneity and tumor microenvironment involvement. (A) Map of the deconvoluted TCGA breast cancer sample proportions of epithelial 1 (Basal), epithelial 3 (HER2) and epithelial 4 (Luminal). Proportions of all three profiles are normalized to equal 100. Dot color indicates the TCGA defined PAM50 subtype. The background color indicates the enrichment of the adipocyte stromal profile vs the CAF stromal profile. Blue indicates a higher proportion of CAFs and yellow represents a higher proportion of adipocytes. (B) Map of the deconvoluted TCGA breast cancer sample. In this figure the background color indicates the proportion of macrophages present in these tumor samples. (C) Histogram showing the correlation between adipocyte markers in the adipocyte stromal profile (yellow) and OXPHOS genes in the epithelial profile, and CAF markers in the CAF stromal profile (blue) and OXPHOS genes in the epithelial profile (blue) (D.1) Map visualizing single cell data and pooled samples across several distinct tumors. The colors of the dots indicate the tumor of origin. In red is sample BC-11, which is then zoomed in on in the next figure. (D.2) Map of single cell sample BC-11(GSE75688, Triple Negative) to visualize the single cell and pooled data for this sample. The pooled sample is comprised of ∼1X10^5^ cells.

There are three possible scenarios that could account for the variation in the location of the tumors within the simplex map: (1) The cancer cells within any specific tumor are relatively pure representatives of Luminal, Basal and HER2 subtypes and that the map position reflects their proportions within the tumor; (2) The cancer cells within a tumor are mostly homogeneous and the position reflects their similarity to the three subtypes; (3) A hybrid between the first two scenarios.

To investigate these possibilities, we leveraged a publicly available single cell RNA-seq dataset (GSE75688) that includes single cell RNA sequencing for breast cancer patients, as well as pooled single cell sequencing and bulk sequencing (Chung et al., 2017). The single cell and bulk RNA-seq was deconvoluted with XDec, and the distribution of the cancer cell profiles was visualized with the simplex map (Figure 3D.1, Figure 3D.2, Figure S3A-H). The single cell and pooled profiles from the same sample did not cluster tightly (Figure 3D.1), suggesting intra-tumoral heterogeneity of cancer cell states. Interestingly, most of the heterogeneity occurred along the Luminal-Basal or Luminal-HER2 axes (Figure 3D.2, Figure S3A-H). This suggests that intra-tumoral heterogeneity partially contributes to the intermediate epithelial profiles seen in bulk tumors in Figure 3A. This conclusion is supported by previous scRNA seq studies which have shown the wide range of heterogeneity in breast cancer tumors (Kim et al., 2018) Taken together, our results suggest that XDec is a useful framework for interpreting single cell RNA-seq data and the epigenetic heterogeneity of cancer cells within individual tumors.

### Mapping reveals heterotypic interactions between cancer cells and the tumor microenvironment

We next investigated interactions between stromal and cancer cells by correlating the transition from adipocyte to CAF stroma with changes within the cancer cells. CAFs have been previously shown to be associated with progression and response to therapy (Brechbuhl et al., 2017; Cazet et al., 2018; Eiro et al., 2019; Hu et al., 2018; Mao et al., 2013; Orimo et al., 2005; Plava et al., 2019; Shiga et al., 2015). Along the Luminal-HER2 axis we observed an increase in the proportion of CAFs (correlation = 0.19) as the proportion of the HER2 profile increased (Figure 3A). We hypothesized that as the cancer cells shift towards the HER2 phenotype, metabolite exchange between cancer cells and the stroma contributes to a more malignant phenotype (Eiro et al., 2019; McCuaig et al., 2017; Plava et al., 2019). There are several molecular mechanisms of metabolite exchange between the epithelial and stromal cells, including exosomes (Boyiadzis and Whiteside, 2015) and other soluble factors (Harper and Sainson, 2014; Popivanova et al., 2009).

Previous studies reported the reverse Warburg effect model in which glycolytic stroma (CAFs) feed lactate and promote oxidative phosphorylation (OXPHOS) in cancer epithelial cells (Martinez-Outschoorn et al., 2015, 2014; Onuchic et al., 2016). These early results were indirect and not based on the deconvolution of the epigenetic states of stromal cells within individual tumors. To correlate the gene expression changes in stromal and epithelial cells directly, we performed Stage 2 deconvolution of the epithelial fractions (STAR Methods). As predicted, we observed a negative correlation between adipocyte gene expression markers in the stromal profile and the expression of epithelial OXPHOS genes. Moreover, the expression of OXPHOS genes in the epithelial fraction correlated positively with stromal CAF markers (Figure 3C). Taken together these correlations suggest metabolic coupling between stromal and cancer epithelial cells consistent with the negative Warburg effect that involves lactate transfer from stroma to cancer epithelial cells.

We also investigated the relationship between cancer cell state, and the immune cell component. While we did not find a correlation between T-cell proportion and cancer cell state, we observed an increase in the proportion of macrophages for the Basal epithelial profile (correlation = 0.22), and a decrease in the proportion of macrophages for the Luminal epithelial profile (correlation = -0.25) (Figure 2B). The increase of Tumor Associated Macrophages (TAMs) was previously associated with Basal breast cancers, non-luminal breast cancers, and predicts poor survival (Gwak et al., 2015; Klingen et al., 2017; Prasmickaite et al., 2018). Thus, the state of the cancer cell is associated with both the stromal and immune profiles of the tumor.

### Matching tumors *in vivo* to PDX models and cell lines

One key challenge in using model systems, such as murine PDX models and cell lines, is to determine which models best represent specific patient tumors. Variability in cell type composition precludes matching by the intrinsic state of the cancer cell based on bulk profiling patient-derived tumor samples.

To explore the correspondence based on the epigenetic state of constituent cancer cells, we projected primary TCGA tumors (left), PDX models (middle), and BRCA cell lines from the Cancer Cell Line Encyclopedia (CCLE) (right) onto the same map (Figure 4A). The density of the distributions indicates that the TCGA tumors are primarily Luminal, the PDX models largely Basal, while the CCLE cell lines show a moderate bias toward the HER2 phenotype. The map therefore provides information about model-specific biases toward specific tumor subtypes while enabling selection of models that best match specific tumors.

**Figure 4.**
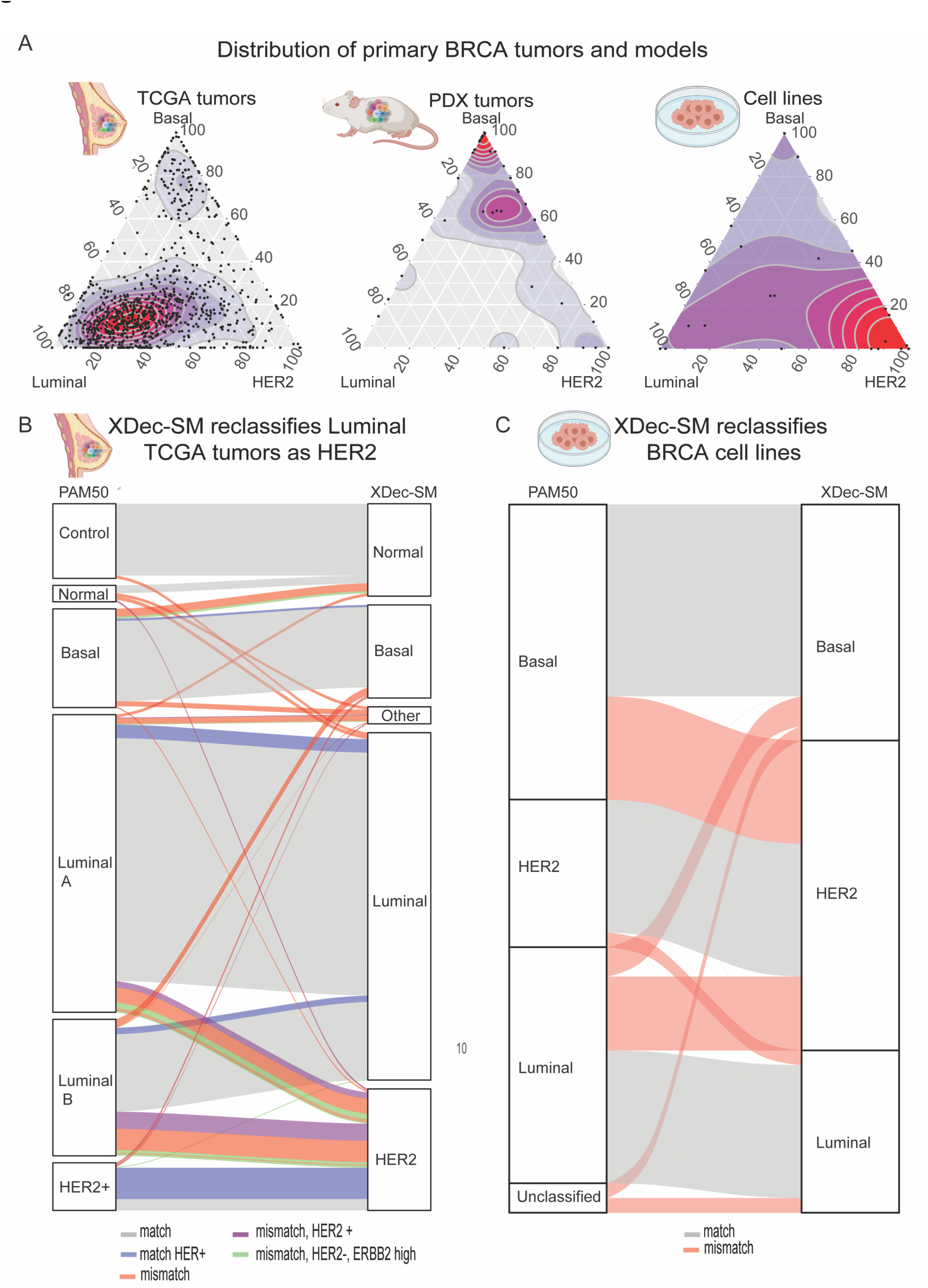
Distribution of samples and classification of cell lines by PAM50 and XDec-SM. (A) Density map of sample position for TCGA (left) PDX (center) and cell lines (right). The density is represented as less dense (blue) to denser (red). The distribution of different sample types varies with PDXs primarily representing Basal tumors and TCGA and cell lines being more widely spread in the simplex. (B) Alluvial plot of TCGA breast cancer sample PAM50 and XDec-SM classification. PAM50 subtypes are provided by TCGA sample metadata. XDec-SM classification indicates the classification based on the maximum proportion of the deconvoluted epithelial profiles. Gray line indicates matching subtype, blue line indicates matching subtype and HER2+ based on the metadata provided by TCGA. Red line indicates mismatching subtype and purple indicates mismatch and HER2+. Green line indicates a mismatch that is HER2--but has a high activation of the ErBB2 pathway. (C) Alluvial plot of XDec-SM clustering classification vs highest proportion for CCLE cell lines. The grey lines indicate samples that were consistently classified by both methods. The red lines indicate samples that are differently classified by both methods.

### Cancer cell state map refines PAM50 classification based on cancer cell-intrinsic states

The PAM50 classification system has been widely utilized for identifying the “intrinsic” subtype of breast tumors. The classification is actionable, with endocrine therapy recommended as the first-line treatment for the tumors classified as Luminal by PAM50 (Marti and Sánchez-Méndez, 2020). In a significant fraction of cases, however, PAM50 classification of tumors does not match HER2 IHC status, another clinically established marker (Raj-Kumar et al., 2019). Notably, PAM50 classification was originally developed based on microarray-based gene expression profiling. In contrast, the map is constructed based on the more informative RNA-seq profiling information. Moreover, PAM50 classification is confounded by the sample-to-sample variation in cell type composition. In contrast, XDec-SM deconvolution provides information about the epigenetic state intrinsic to the cancer cell fraction of the tumor. We therefore asked how the two classifications compared in primary TCGA breast cancer tumors. For each tumor we obtained PAM50 classification assigned in the TCGA database and determined XDec classification by identifying the cancer profile (Luminal, HER2, Basal) with the highest proportion (closest corner in the map). Most tumors had the same classification (Figure 4B, grey line). As expected, HER2+ tumors (Figure 4B, blue line) were primarily classified as HER2 by both methods. The largest discrepancies involved tumors classified as Luminal by PAM50 and HER2 by XDec-SM (Figure 4B, red line). A subset of those tumors was HER2 positive (Figure 4B, purple), or had a high ErBB2 signaling network activation score developed by Lim et al. (Figure 4B, green line), concordant with XDec-SM classification, suggesting that the state at least some of the cases reflect ErBB2 pathway activation. We then proceeded to ask if we could also use the XDec-SM classification method to characterize breast cancer cell lines.

Cancer cell lines are an important model system, and datasets such as the CCLE have characterized large numbers of them. Because cell lines often evolve *in vitro* and differ from primary tumors, there is long-standing interest in identifying the cell lines that best model specific types of breast tumors (Jiang et al., 2016). PAM50 classification has been utilized for both cell lines and tumors, thus helping match the tumors and cell lines by type. However, there are several drawbacks to applying PAM50 to cell lines. Primarily, PAM50 was developed on bulk array data, and contains stromal genes that are not intrinsically represented in cell lines. We focused on characterizing the state of cell lines by placing them on the same cancer cell state map as deconvoluted primary tumors.

To characterize commonly used breast cancer cell lines, we obtained the expression data from 51 cell lines profiled by the CCLE (Ghandi et al., 2019). The PAM50 annotations for the cell lines were obtained from the study by Jiang et al (Jiang et al., 2016). We classified the cell lines by their map position, by assigning them to their closest vertex (Figure 4C). As with the primary TCGA breast cancer tumors, for the majority of cell lines the PAM50 and XDec-SM classification were concordant. However, a fraction of cell lines that were classified as Luminal or Basal by PAM50 were reclassified as HER2 by the XDec-SM classification method. As the XDec-SM classification method is based on cancer cell state, we were also able to classify cell lines which were previously unclassified by the PAM50 method (Figure 4C).

### Cancer cell state associates with driver mutations, key pathway activations, and DNA methylation patterns

We next asked how the positions on the map correlate with mutations in key tumor suppressors, oncogenes, and with the activation of pathways driving cancer progression. As indicated in Figure 5A, mutations in PIK3CA localize in the Luminal corner, while TP53 mutations localize in the Basal corner consistent with the high subtype-specific prevalence of mutations in these genes. In contrast TNN mutations do not show subtype preference.

**Figure 5.**
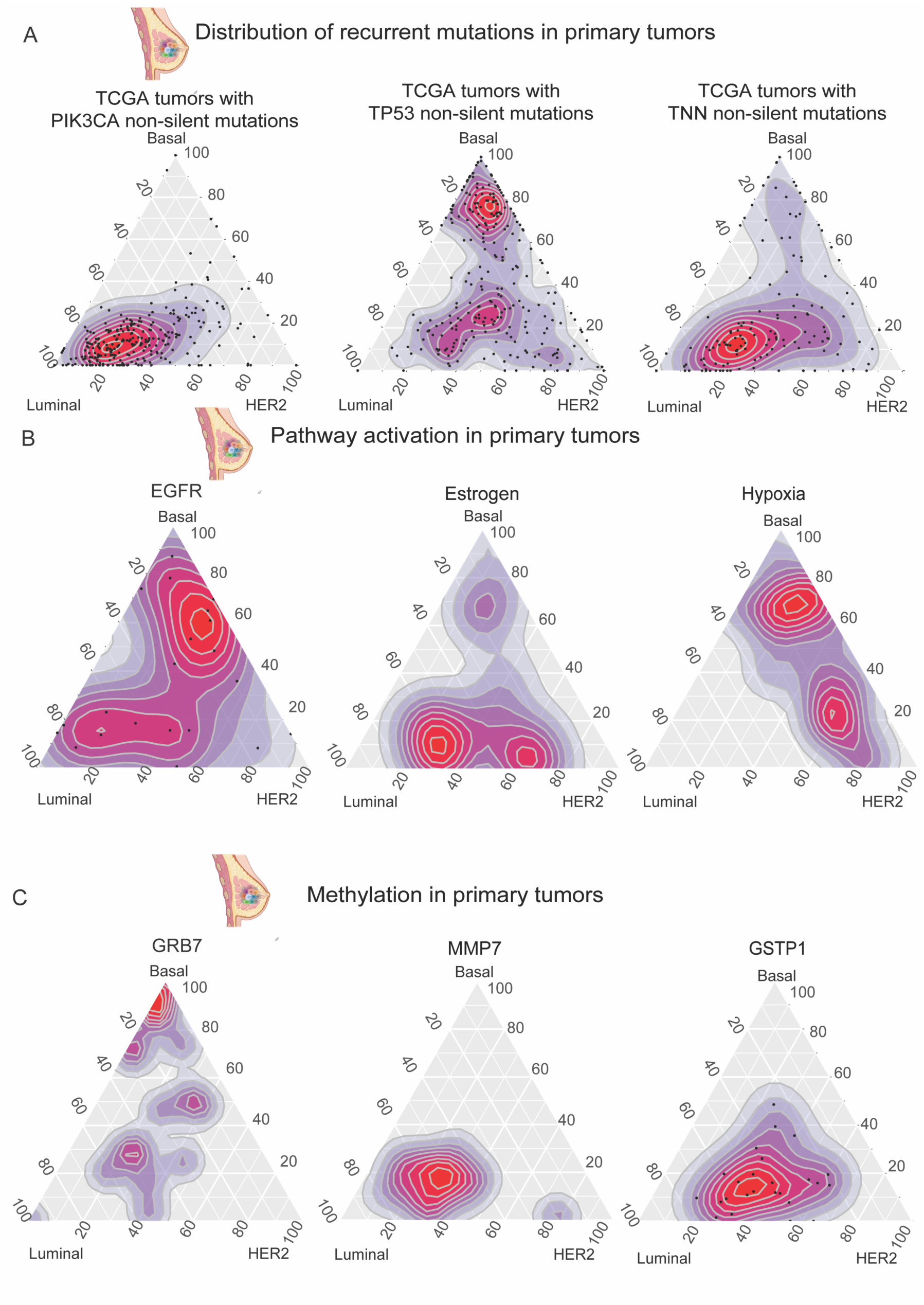
Distribution of mutations, pathway activation, and methylation profiles in cancer cell state map. (A) Distribution of common breast cancer mutations including PIK3CA (left), TP53 (middle), and TNN (right). The density is represented as less dense (blue) to denser (red). (B) The activation of MAPK, P53, Hypoxia, and NFk-B activation as projected on the map. The tumors with the 25 highest activation scores are plotted, with the density is represented as less dense (blue) to denser (red). (C) Methylation patterns of GRB7, MMP7, and GSTP1. The 25 tumors with the highest average methylation scores across all loci in each gene are plotted on the cancer cell state map. Density is less dense (blue) to most dense (red).

We then used the cancer cell state map as a framework to visualize pathway activation by plotting the 25 tumors with the highest pathway activation score for EGFR, estrogen, and hypoxia. EGFR pathway activation is more activated in the Basal subtype, with the estrogen pathway being active in Luminal and HER2 subtypes (Figure 5B). Basal tumors show highest activation for Hypoxia, consistent with inactivating TP53 mutations protecting the Basal cancer cells from the apoptotic effects of the TP53 pathway.

The cancer cell state was also used to visualize the methylation profiles of the TCGA breast cancer tumors (Figure 5C). As with the pathway analysis, the tumors that had the highest level of methylation (n = 25) were plotted on the map for GRB7, MMP7, and GSTP1. In concordance with previous studies (Holm et al., 2010), GRB7 is the most methylated in the Basal corner. Likewise, MMP7 and GSTP1 which have been associated with luminal A and luminal B subtypes respectively are found to be the most methylated along the Luminal to HER2 axis. In summary, these mutational, pathways activation, and methylation patterns recapitulate known breast cancer biology, suggesting that the position in the map should be highly informative for the state and behavior of the cancer cell.

### Cancer cell state in PDX models is predictive of chemotherapy response

We next asked if the epigenetic state of cancer cells in PDX models was predictive for chemotherapy response. Forty-four PDX models from the BCM collection with known response to carboplatin and docetaxel were deconvoluted. Information about these PDX models can be accessed through the PDX portal (https://pdxportal.research.bcm.edu/). The Basal, Luminal, and HER2 proportions of these PDXs were used to build generalized linear models (GLMs) of response to both therapies.

The GLM models built on the BCM PDX cohort were used to predict the chemotherapy response of an independent dataset consisting of 22 PDXs from the Rosalind & Morris Goodman Cancer Research Centre (RMGCRC) (GSE142767) (Savage et al., 2020). The response to both agents in the RMGCRC cohort was annotated with RECIST criteria as complete response (CR), partial response (PR), stable disease (SD), or progressive disease (PD). While the training and test datasets did not use the same platinum and taxane compounds, they were matched by compound class (carboplatin GLM used to predict cisplatin response and docetaxel GLM used to predict paclitaxel response). With only the three predictor variables (Luminal, Basal, HER2 proportion), we found that the GLM models had high predictive power for the platinum agents (ROC complete response vs all else (CR) = 0.73, ROC complete and partial response vs all else (CRPR) = 0.80) and to a lesser degree the taxanes (ROC complete response vs all else = 0.58 (CR), ROC complete and partial response vs all else (CRPR) = 0.67) (Figure 6A). Taken together, these results suggest that the state of cancer cells in PDX models is predictive of response to both platinum and taxane therapies. A similar approach could be utilized to predict the response of patients to neoadjuvant chemotherapy.

**Figure 6.**
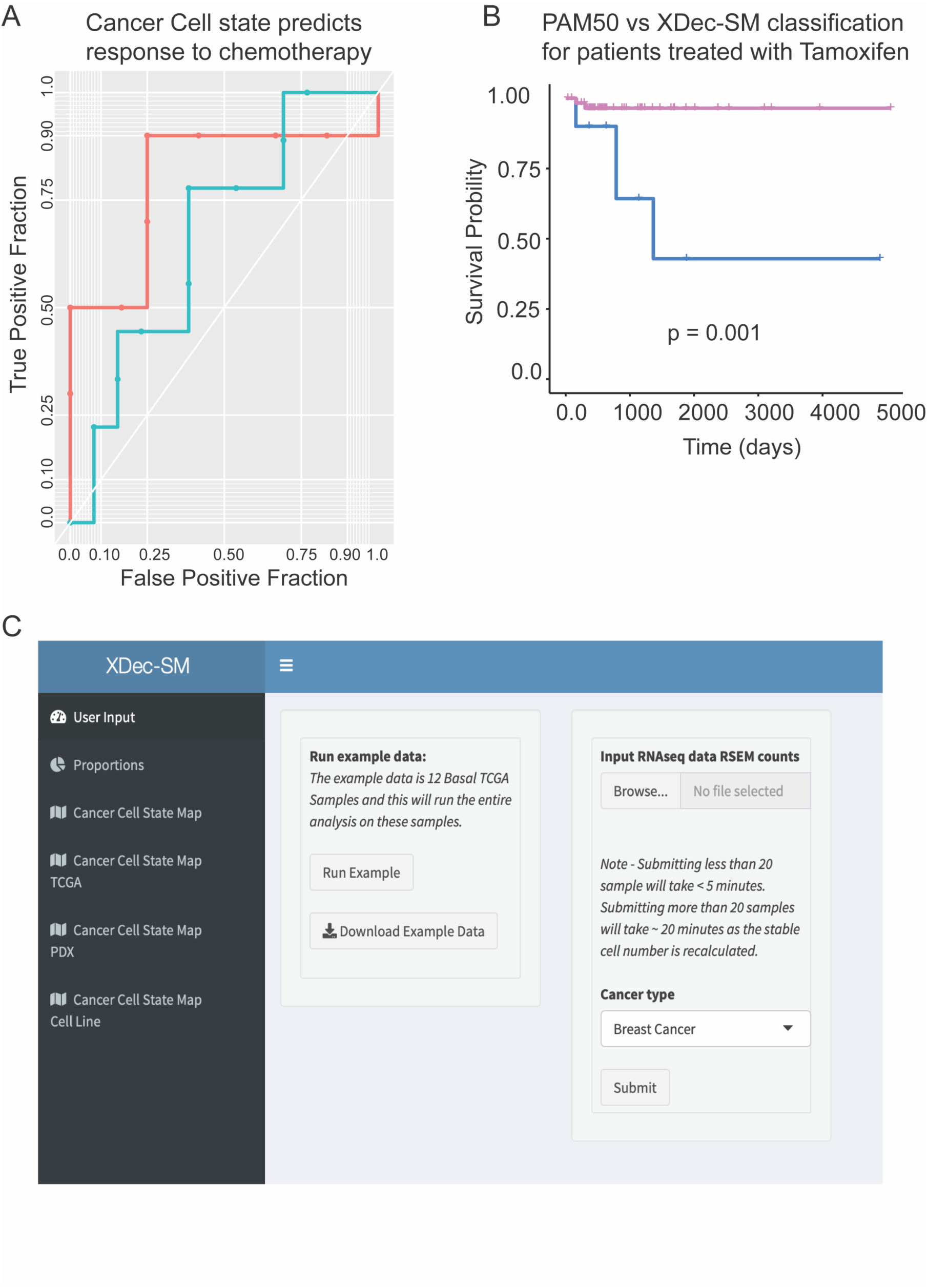
Cancer cell state map is predictive of therapy response. (A) ROC curve of the response to cisplatin and paclitaxel in the Goodman PDX dataset as predicted using a GLM model built on the epithelial proportions and the carboplatin and docetaxel response in the BCM PDX dataset. CRPR vs all else for platinum agents is in red (with an AUC of 0.80) and CRPR vs all else for taxanes is in blue (with an AUC of 0.67). (B) Kaplan-Meier Curve that indicates the difference in survival between samples classified as Luminal by PAM50 and XDec-SM (pink) and samples classified as Luminal by PAM50 and HER2 by XDec-SM (blue). (C) Online deconvolution tool. Screenshot of the XDec-SM deconvolution tool available at https://brl-bcm.shinyapps.io/XDec_BRCA/.

### Cancer cell states in patient tumors predicts response to tamoxifen

As the state of the cancer cell is predictive of response to chemotherapy, we then asked if the XDec-SM classification itself was predictive of response. In the previous section titled “*Cancer cell state map refines PAM50 classification based on cancer cell-intrinsic states*” we found that the largest discrepancy between the PAM50 and XDec-SM classification of TGCA tumors was that a subset of tumors that were classified as Luminal by PAM50 were classified as HER2 by XDec-SM. Additionally, some of tumors that were reclassified as HER2 by XDec-SM showed ErBB2 pathway activation which is a negative predictor of tamoxifen response.

To examine whether cancer cell state indeed identifies cancer cells that respond poorly to tamoxifen, we next focused on 72 TCGA BRCA tumors that were classified as Luminal by PAM50 (using recount package (Collado-Torres et al., 2017)) treated with tamoxifen, and had survival information (NCI GDC annotations). Of those, 62 were classified as Luminal and 10 as HER2 by XDec-SM. The two groups showed survival differences, with patients classified as HER2 by XDec-SM alone having significantly worse survival (Figure 6B, p = 0.001). Notably, neither HER2 status alone (Figure S4A), nor ErbB2 score (Figure S4B), could identify subsets of patients with significantly worse survival (p=0.4 and 0.77 respectively). This suggests that cancer state alone could identify a subset of poor responders to tamoxifen therapy among those that were classified as Luminal by PAM50.

## DISCUSSION

To construct the cancer cell state map of breast cancer, we developed the XDec-SM deconvolution algorithm. Unlike other RNA-seq based deconvolution methods that are based on reference profiles known *a priori* from profiling of previously isolated cell populations, XDec-SM is reference-optional, thus enabling data-driven discovery of novel epigenetic states of cells within tumors and other complex tissues. Remarkably, the algorithm produces data-driven estimates of constituent cell types that correspond to PAM50 (Nielsen et al., 2010; Parker et al., 2009) and the current molecular classification of breast cancer into Luminal, HER2 and Basal subtypes. Unlike other similar methods (Rolong et al., 2021), XDec-SM allows us to map tumors onto a low dimensional (2D) simplex map. Importantly, this simplex map is biologically interpretable, with each vertex anchored at a known breast cancer subtype. By mapping breast cancer tumors onto this simplex map, we refine the PAM50 classification by placing tumors on the spectrum between established PAM50 subtypes.

Additionally, unlike other RNA-seq based deconvolution methods that focus on explaining variance in levels of RNA, XDec-SM in Stage 1 scales gene expression levels within the 0-1 range, thus focusing on the information about the epigenetically regulated on/off states of gene transcription. The epigenetic state of the cell is programmable during development and intrinsic factors such as hormonal signaling, paracrine signaling and other heterotypic interactions between cells within complex tumor tissue leave an “epigenetic footprint”. The epigenetic state of the cell can in principle be estimated by measuring either “classical” epigenetic marks such as DNA methylation or gene expression levels. However, molecular array or sequencing-based profiling of bulk tumor tissue is confounded by the sample-to-sample variability in cell type composition, precluding access to the intrinsic epigenetic states of constituent cell types. By deconvoluting the cell state with XDec-SM, we can capture the cancer cell states in the sample and obtain a snapshot of the “epigenetic footprint” of the tumor projected onto the cancer cell state map. In this manner, XDec-SM extends the previously proposed Epigenomic Deconvolution method based on bulk DNA methylation profiles demonstrating that the epigenetic heterogeneity of tumors can be concordantly assessed from bulk RNA-seq or DNA methylation.

While XDec-SM benefits from information about which genes are informative for deconvolution gleaned from publicly accessible scRNA-seq profiles of tumor cells, it is also designed to infer new recurrent states of cancer cells not obvious from the analysis of scRNA-seq data. Thus, allowing the method to be reference-optional and leverage information from both the informative cell type specific loci and PAM50 literature-based gene signatures. The cancer cell state map is synergistic and complementary with these scRNA-seq methods, as it starts from bulk RNA-seq profiles and thus provides an embedding based on vastly more samples, including those without single-cell data covering a larger diversity of tumors albeit at lower resolution. Because bulk RNA-sequencing data is available for many more samples and does not involve perturbation of the cell state associated with physical isolation XDec-SM approach is uniquely positioned to provide a view of the full spectrum of recurrent states of cancer cells *in vivo*.

We show that the cancer cell state map of breast cancer not only extends established tumor classification, but also provides additional information about tumor biology, model systems, and drug response. We demonstrate that the cancer cell state map is also useful as a framework for interpreting scRNA-seq profiles of individual tumors: by placing scRNA-seq profiles on the map, we identify significant epigenetic heterogeneity within individual tumors, particularly along the Luminal-Basal and Luminal-HER2 axes.

Our findings demonstrate that cancer cell state map helps visualize and quantitatively analyze (by map position) heterotypic interactions within tumors. By correlating the states of stromal and cancer cell fractions, we recapitulate metabolic coupling between them, consistent with the previously reported reverse Warburg effect (Onuchic et al., 2016). There are several mechanisms through which metabolite exchange between the epithelial and stromal cells can occur, including soluble factors (Harper and Sainson, 2014; Popivanova et al., 2009) and exosomes (Boyiadzis and Whiteside, 2015). More broadly, our findings reinforce the role of cancer-associated fibroblasts in cancer (Brechbuhl et al., 2017; Cazet et al., 2018; Eiro et al., 2019; Hu et al., 2018; Mao et al., 2013; Orimo et al., 2005; Plava et al., 2019; Shiga et al., 2015). Additionally, the map sheds light on the association of tumor-associated macrophages with the Basal subtype. Because some of the heterotypic interactions may be mediated by exosomes and other highly specific factors that may be found in bodily fluids, the cell-cell communication discovered by cell state mapping may open the doors toward the development of liquid biopsy biomarkers.

Additionally, the precise characterization of the state of the cancer cell fraction within individual tumors improves the matching of individual tumors with specific experimentally accessible PDX and cell line models. The map places breast cancer cell lines on the epigenetic spectrum between established breast cancer subtypes, identifying the subsets that can serve as subtype-specific models, versus those that do not clearly model any established subtype of breast cancer. This information is critical for understanding the translational potential of *in vitro* therapeutic drug screens (Ghandi et al., 2019) as each breast cancer subtype may have a distinct response (Holliday and Speirs, 2011). We demonstrate the utility of this low dimensional to map to predict therapy response in experimentally tractable models and tumors *in vivo*. The map position predicts response to standard chemotherapies based on PDX models and targeted therapies based on cell line models. Cancer cell state mapping identifies a subset of tumors classified as Luminal by PAM50 (Nielsen et al., 2010; Parker et al., 2009) that are HER2-like and are resistant to tamoxifen therapy, a first-line therapy for this predominant subtype of breast cancer. While these preliminary results are based on a small number of samples, they demonstrate the potential utility of the application of cell state mapping in precision medicine.

We note that the XDec-SM algorithm is very generic and is not limited to deconvolution of solid tissues. Specifically, XDec-SM has previously been applied to deconvolute thousands of extracellular RNA-seq profiles of human body fluids in the exRNA Atlas and to construct the first reference map of extracellular RNA in human body fluids (Murillo et al., 2019). Analogously to the scRNA-seq profiles, the selection of informative features was informed by publicly available RNA-seq profiles from experimentally isolated exRNA carriers (vesicle, lipoprotein, RNA-binding protein).

To empower the community to use this method, we made the XDec-SM code available online (Figure 6C) and as a R package under a free open-source license (https://github.com/BRL-BCM/XDec). We also developed an interactive web service that allows users to deconvolute breast tumor RNA-seq profiles of interest and project them onto the cancer cell state map (https://brl-bcm.shinyapps.io/XDec_BRCA/).

## LIMITATIONS OF THE STUDY

The power of XDec-SM method is yet to be fully characterized and it may be hard to characterize under realistic assumptions. For example, while several states of breast cancer cells were identified, is not clear how many additional profiles would be required to identify the diverse states within stromal cell types. The granularity of the XDec-SM deconvolution is also dependent of the availability of single cell reference profiles and the amount of sample-to-sample variability in cell type proportions and the variability in the proportions of cells in specific states. While the proliferation of single cell studies and bulk profiling will help increase the resolution of the map over time, because of the interplay of relevant parameters, it would be hard to estimate, under realistic assumptions. the number and types of profiles required for such high-resolution map.

## ACKNOWLEDGEMENTS

This work was supported by a grant from the Common Fund of the National Institutes of Health (NIH) (5U54 DA036134) (to A.M.), an NCI PDX Development and Trials Center grant U54CA224076 (to M.T.L), a CPRIT Core Facility Support Grant RP170691 (to M.T.L.) and P30 Cancer Center Support Grant (NCI-CA125123).

## AUTHOR CONTRIBUTIONS

Conceptualization, O.D.M., V.P, M.T.L, A.M.; Methodology, O.D.M., V.P., E.L.L., L.E.D, and A.M.; Software, O.D.M., V.P., and E.L.L.; Validation, O.D.M., and V.P.; Formal Analysis, O.D.M., V.P., and A.M.; Investigation, O.D.M., V.P., E.L.L, and A.M.; Resources, L.E.D, M.T.L, A.M.; Data Curation, O.D.M., V.P., and E.L.L; Writing O.D.M., V.P, and A.M.; Visualization, O.D.M., V.P., and A.M.; Supervision, A.M.; Project Administration, A.M.; Funding Acquisition, M.T.L, and A.M.

## STAR METHODS

### RESOURCE AVAILABILITY

#### Lead Contact

Further information and requests for resources should be directed to and will be fulfilled by the Lead Contact, Aleksandar Milosavljevic.

#### Material Availability

This study did not generate new unique reagents.

#### Data And Code Availability

The datasets generated during this study are available on Genboree Commons (http://genboree.org/theCommons/projects/xdec-singlecell) and code is available on Github (https://github.com/BRL-BCM/XDec).

### METHOD DETAILS

#### XDec-SM Deconvolution Algorithm

XDec-SM is a variation of the EDec algorithm, adjusted for RNA-seq data. EDec is an iterative algorithm for constrained matrix factorization using quadratic programming for DNA methylation-based deconvolution (Onuchic et al., 2016). The key difference is that in Stage 2 of XDec-SM expression values are transformed by a single-parameter negative exponential function into the 0-1 range (the same range that applies to original DNA methylation). XDec-SM has originally been applied to deconvolute extracellular RNA-seq profiles of human bodily fluids (Murillo et al., 2019). Like EDec, XDec-SM has three stages: Stage 0, 1, 2 (Figure 1A).

XDec-SM relies on a set of informative features that are differentially abundant across each component modeled in the system (Figure 1A, Stage 0). The set of informative genes may be obtained by RNA sequencing of cell lines or by single-cell RNA sequencing. In case of exRNA deconvolution, this information is provided by the RNA sequencing of the cargo of specific carriers of extracellular RNA in human body fluids (various vesicles, lipoprotein particles, and RNA-binding proteins). In any case these profiles are used for the sole purpose of identifying sets of informative loci and otherwise do not bias the output of deconvolution.

Stage 1 involves iterative matrix factorization to estimate the gene expression profiles of constituent cell types and the per-sample proportions of constituent cell types in individual tumors. As mentioned above, for the purpose of this computation, expression counts are transformed to the [0-1] range using the following negative exponential function:

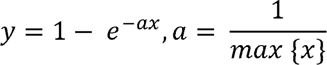

In Stage 2 (Figure 1A, Stage 2), XDec-SM uses the per-sample proportions estimated in Stage 1 to estimate the untransformed gene expression profile of each constituent cell type utilizing constrained least-squares fit using quadratic programming. This gene expression profile represents the average gene expression across the set of samples.

#### Cell Line RNA Sequencing Data Processing

This set represented 7 cell types: epithelial (GSM1695870, GSM1695871, GSM1695872, GSM1695873, GSM1695874, GSM1695875, GSM1695876, GSM1695877, GSM1695878, GSM1695879, GSM1401648, GSM1401649, GSM1401650, GSM1401651, GSM1401653, GSM1401654, GSM1401665, GSM1401668, GSM1401672, GSM1401673, GSM1401674, GSM1401675), B cells (GSM1576391, GSM1576419), dendritic cells (GSM1576395, GSM1576423), monocytes (GSM1576399, GSM1576427), CD56+ natural killer cells (GSM1576407, GSM1576435), T cell (GSM1576415, GSM1576443) and stroma (GSM2430225, GSM2430226, GSM2430227, GSM2839374). Transcript fastq files were downloaded from GEO and then aligned and quantified using STAR (Version 2.4) and RSEM (Version 1.3). The reference genome was assembled using the standard rsem-prepare-reference function using the hg19 genome assembly. RSEM was run using standard parameters (rsem-calculate-expression –star -p 6 –star-gzipped-read-file –paired-end) to produce RSEM gene counts for each of the cell line profiles. The resulting 36 profiles all met high read quality standards using fastqc with standard parameters. RSEM gene expression profiles for all 36 profiles were transformed using Equation 1 to constrain the values to a [0-1] range. Genes with low coverage (=< 0.01) and those not identified within the TCGA breast cancer profiles were removed from further analysis.

#### scRNA Sequencing Data Processing

Single cell gene expression RSEM counts (GSE118389_counts_rsem) were downloaded from GEO (GSE118389). Metadata provided by the study determined the cell type identity (Karaayvaz et al., 2018). For XDec-SM, endothelial and B cell profiles were removed from further analysis due to low number of profiles. Initially all single cell profiles per cell type are ranked based on total read coverage. Profiles with low coverage across all genes are moved (sum of gene counts < 100,00) are removed. Every five profiles are then summed to create pseudo bulks (Lun et al., 2016). Gene reads for each pseudo bulk are then normalized to match the highest coverage pseudo bulks across all cell types. All pseudo bulks are transformed using Equation 1 to a 0-1 range. This results in pseudo bulks for four cell types: epithelial (n = 128), stromal (n = 18), T cell (n = 10), macrophages (n = 12).

#### TCGA Data Processing

The TCGA BRCA dataset (IlluminaHiSeq_RNASeqV2) and clinical phenotype metadata (BRCA_clinicalMatrix) were downloaded from UCSC Xena (https://xena.ucsc.edu). Only samples with matching gene expression profiles and metadata (n = 1,218) were kept for further analysis. PAM50_mRNA_nature2012 metadata information was used for the subtype determination. Samples with “-11” were designated as healthy controls. TCGA samples were quantile normalized and transformed using Equation 1 to constrain gene expression values to a 0-1 range. Genes with low gene expression coverage across all samples (average < 0.01) were removed.

### QUANTIFICATION AND STATISTICAL ANALYSIS

#### XDec-SM Application to Simulated Dataset

To test the XDec-SM methodology, we first simulated 100 mixtures based on publicly available, experimentally isolated gene expression profiles (twelve breast cancer epithelial, ten normal epithelial, ten immune, and four stromal, all listed below). The gene expression profiles for each cell type were randomly chosen and transformed into the [0-1] range. These profiles were then mixed in varying proportions, and random noise was added to simulate heterogenous tumor data (described below).

Stage 0 was performed to identify differentially expressed genes (n = 226) across the reference profiles of the 4 cell types. XDec-SM Stage 1 was applied to the simulated mixture samples and 4 constituent cell types were modeled. The estimated expression profiles are correlated to the reference expression profiles over the informative gene set to ascertain the corresponding identity of each cell type. XDec-SM can accurately deconvolute the gene expression profile of each cell type when compared to the reference profile with the greatest correlation (cancer epithelial, R^2^ = 0.88; normal epithelial, R^2^ = 0.97; immune, R^2^ = 0.96; stromal, R^2^ = 0.98). Additionally, XDec-SM accurately estimates the per-sample proportion of all four cell types (cancer epithelial, R^2^ = 0.98; normal epithelial, R^2^ = 0.97; immune, R^2^ = 0.96; stromal, R^2^ = 0.96). Thus, both the simulated data was used to validate both the transformation procedure and the XDec-SM methodology.

To generate the simulated mixture samples, a set of 36 cell line expression profiles collected from GEO (https://www.ncbi.nlm.nih.gov/geo/) were utilized. This set represented 4 major cell types: normal epithelial (GSM1695870, GSM1695871, GSM1695872, GSM1695873, GSM1695874, GSM1695875, GSM1695876, GSM1695877, GSM1695878, GSM1695879), cancer epithelial (GSM1401648, GSM1401649, GSM1401650, GSM1401651, GSM1401653, GSM1401654, GSM1401665, GSM1401668, GSM1401672, GSM1401673, GSM1401674, GSM1401675), immune (GSM1576391, GSM1576395, GSM1576399, GSM1576407, GSM1576415, GSM1576419, GSM1576423, GSM1576427, GSM1576435, GSM1576443), and stroma (GSM2430225, GSM2430226, GSM2430227, GSM2839374). The cancer epithelial cell lines include a diverse set of breast cancer cell lines of varying subtypes (ZR-75-1, BT-474, MDA-MB-468, HCC38, MCF-7, T-47D, HCC1954, HCC1937, HCC1187, BT-20, HCC1569, SUM-102). Immune cell lines include two profiles for each subtype: B cells, dendritic cells, monocytes, CD56+ natural killer cells, and T cells. Stromal cell lines represent cancer associated fibroblasts (CAFs). Transcript fastq files were downloaded from GEO and aligned and quantified using STAR (Version 2.4) and RSEM (Version 1.3). The reference genome was assembled using the standard rsem-prepare-reference function using the hg19 genome assembly. RSEM was run using standard parameters (rsem-calculate-expression --star -p 6 --star-gzipped-read-file --paired-end) to produce RSEM gene counts for each of the cell line profiles. The resulting 36 profiles all met high read quality standards using fastqc with standard parameters (https://github.com/s-andrews/FastQC). RSEM gene expression profiles for all 36 profiles were transformed as described above to constrain the values to the [0-1] range. Genes with low coverage (=< 0.01) are removed from further analysis.

An independent gene expression random variable was used to generate noisy versions of the expression profiles. The random variable was selected as 10% of the maximum variance for the cancerous epithelial profiles and 5% for the normal cell types (normal epithelial, stromal, immune). Once the noisy profiles were generated, each simulated mixture contains a randomly selected profile for each of the 4 cell types. The simulated mixtures are a linear combination of the noisy expression profiles and a set of proportions for each of the 4 cell types. High purity mixtures included greater than 60% cancer epithelial, impure samples include 40% cancer epithelial, low purity included 10% cancer epithelial and over 50% normal epithelial, and control samples included 0% cancer and 70% normal epithelial. The remaining proportions are divided by the stromal and immune cell types.

Using the 4 cell type classes (cancerous epithelial, normal epithelial, immune, stroma), XDec-SM Stage 0 was performed to identify informative genes. We performed t-test across the 4 cell type classes comparing the transformed gene expression values between each group of reference profiles of the same cell type to the remaining expression profiles. For those genes with a significant differential expression (p.value < 0.0001), the 25 most upregulated and the 25 most downregulated genes were selected to represent each cell type. To further separate cancerous epithelial and stroma and normal epithelial and stroma, a direct t-test comparison was performed comparing the each of the two groups. For those genes with a significant differential expression (p.value < 0.0001), the 25 most upregulated and the 25 most downregulated genes were selected. If any genes appeared multiple times, they were excluded from the list of informative genes resulting in 226 genes.

XDec-SM Stage 1 was performed using the 100 simulated mixtures as input (4 normal epithelial, 4 immune and 4 stromal profiles were included in the input datasets for stability). Deconvolution was performed using the 226 informative genes, a preset number of cells (n = 4), max iterations = 2000, and residual sum of squares differential stop = 1e-10.

#### XDec-SM Application to TCGA Dataset with Informative Genes from Cell Line Profiles

Using the 7 cell type classes (epithelial, B cells, dendritic cells, monocytes, CD56+ natural killer cells, stroma), XDec-SM Stage 0 was performed to identify informative genes. We performed t-test across the 7 cell type classes comparing the transformed gene expression values between each group of reference profiles of the same cell type to the remaining expression profiles. For those genes with a significant differential expression (p.value < 0.0001), the 25 most upregulated and the 25 most downregulated genes were selected to represent each cell type. To further separate epithelial and stroma, a direct t-test comparison was performed comparing the two groups. For those genes with a significant differential expression (p.value < 0.00001), the 75 most upregulated and the 75 most downregulated genes were selected. If any genes appeared multiple times, they were excluded from the list of informative genes resulting in 391 genes. The PAM50 gene set was also included to define subtype specific expression for a total of 440 informative genes (49 out of the 50 PAM50 genes were identified in the TCGA dataset).

XDec-SM Stage 1 was performed using the 1,215 TCGA bulk tumor samples as input (4 normal epithelial, 5 immune and 4 stromal profiles were included in the input datasets for stability). Deconvolution was performed using the 440 informative genes, max iterations = 2000, and residual sum of squares differential stop = 1e-10. Stability criteria was used to identify a stable number of cells in the model. Using 3 replicates of 80% of the input data, deconvolution is performed modeling 3 to 10 cell types. The estimated proportions from the three replicates are compared for each cell type model and the number of cell types is determined once the correlations are no longer significant. Stability for this model resulted in 6 cell types.

#### XDec-SM Application to TCGA Dataset with Informative Genes from Single-Cell Profiles

XDec-SM Stage 0 was performed to identify differentially expressed genes across the pseudo bulk profiles of the 4 cell types (epithelial, stromal, T cells, macrophages), XDec-SM Stage 0 was performed to identify informative genes. We performed t-test across the 4 cell type classes comparing the transformed gene expression values between each group of pseudo profiles of the same cell type to the remaining expression profiles. For those genes with a significant differential expression (p.value < 0.05), the 50 most upregulated and the 50 most downregulated genes were selected to represent each cell type. To further separate T cells and macrophages, a direct t-test comparison was performed comparing the two groups. For those genes with a significant differential expression (p.value < 0.05), the 25 most upregulated and the 25 most downregulated genes were selected. If any genes appeared multiple times, they were excluded from the list of informative genes resulting in 274 genes. Additionally, we included the PAM50 gene set to identify any subtype specific signatures for a total of 323 informative genes (274 plus 49 out of the 50 PAM50 genes).

XDec-SM Stage 1 was performed using the 1,215 TCGA bulk tumor samples as input. Deconvolution was performed using the 323 informative genes, max iterations = 2000, and residual sum of squares differential stop = 1e-10. Stability criteria was used to identify a stable number of cells in the model. Using 3 replicates of 80% of the input data, deconvolution is performed modeling 3 to 12 cell types. The estimated proportions from the three replicates are compared for each cell type model and the number of cell types is determined once the correlations are no longer significant. Stability for this model resulted in 9 cell types.

XDec-SM Stage 2 was performed for three distinct analyses. First, TCGA breast cancer samples were subset into five groups based on the predominant epithelial proportion (epithelial 1 (Basal) [n = 133], epithelial 2 (Normal Control) [n = 107], epithelial 3 (HER2) [n = 176], epithelial 4 (Luminal) [n = 510], epithelial 5 (Other) [n = 30]). Proportions of the nine deconvoluted profiles were combined into three main cell types: epithelial (epithelial 1 through 5), stromal (stromal 1 and stromal 2), and immune (T cell and macrophage).

Second, XDec-SM Stage 2 was performed on each of the PAM50 breast cancer subtypes. This resulted in 6 cohorts: Controls [n = 95], Normal-like [n = 24], Basal-like [n = 142], Luminal A [n = 422], Luminal B [n = 194], HER2 [n = 67]. The proportions of the two stromal cell profiles (stromal 1 and stromal 2) were not combined in this analysis but the epithelial (epithelial 1 through 5) and immune (T cell, macrophage) cell profiles were combined respectively.

Lastly, XDec-SM Stage 2 was performed on each of the novel breast cancer subtypes with methylation data. This resulted in 4 groups: control [n = 73], Basal [n = 85], HER2 [n = 102], Luminal [n = 321]. The proportions of epithelial, immune, and stromal components were combined in this analysis.

#### Deconvolution Methods

##### CIBERSORTx

Single cell gene expression RSEM counts (GSE118389_counts_rsem) were downloaded from GEO (GSE118389). Metadata provided by the study determined the cell type identity (Karaayvaz et al., 2018). Using the CIBERSORTx web interface (https://cibersortx.stanford.edu), the “Create Signature Matrix” module (default parameters) was used to create the signature matrix for the 6 cell types: epithelial, stroma, endothelial, B cell, T cell, and macrophage. The TCGA BRCA dataset (IlluminaHiSeq_RNASeqV2) was downloaded from UCSC Xena (https://xena.ucsc.edu) and was uploaded as the “Mixture file”. The “Impute Cell Fractions” module was run using default parameters using the previously defined Signature Matrix. For comparison to other methods, the three immune cell fractions (B cell, T cell, macrophage) are summed to represent immune and the two stromal cell fractions (stroma, endothelial) are summed to represent stromal.

##### InfiniumPurify

The TCGA BRCA proportions were previously estimated (Qin et al., 2018) and downloaded from Zenodo (https://zenodo.org/record/253193#.Xr1_xy-z2uV). Only TCGA samples with both XDec-SM proportions and InfiniumPurify tumor purity were compared.

##### TIMER

The TCGA BRCA proportions were previously estimated (Li et al., 2017)and downloaded (http://timer.cistrome.org). Only TCGA samples with both XDec-SM proportions and TIMER immune scores were compared. TIMER estimated the immune score for 6 immune cell subtypes (B_cell, CD4_Tcell, CD8_Tcell, Neutrophil, Macrophage, Dendritic) and all scores were summed for comparison. ***Epigenomic Deconvolution (EDec)***

The TCGA BRCA proportions were previously estimated (Onuchic et al., 2016) and were downloaded from Genboree (http://genboree.org/theCommons/projects/edec). Only TCGA samples with both XDec-SM and EDec estimated proportions were compared. The six epithelial profiles (cancerous epithelial 1-5, normal epithelial) are summed to represent the epithelial cell type proportions.

##### Mapping of Samples onto the Cancer Cell State Map

After the deconvolution of all samples (TCGA, PDXs, cell line, single cell profiles), only samples with a combined epithelial proportion of > 0.7 and an epithelial Basal + epithelial HER2 + epithelial Luminal proportion of > 0.1 were selected to ensure that these samples were well modeled by the three epithelial profiles. The samples were then placed on a cancer cell state map representing their cancer cell state as a combination of the epithelial Basal, epithelial, HER2 and epithelial Luminal profiles. To visualize the simplex, the proportions of the sum of these three profiles were normalized to equal 1. The ggtern package was utilized to visualize the map (Hamilton and Ferry, 2018).

##### Association of Tumor Microenvironment and Cancer Cell Profiles

After the deconvolution of TCGA samples with XDec-SM, a correlation analysis was done to determine the relationship between the proportion of CAFs and the proportion of the epithelial HER2 profile. Likewise, a correlation was done between macrophage proportion and the proportion of the epithelial Basal and epithelial Luminal profiles.

##### Classification of TCGA and Cell Lines

For the TCGA XDec-SM classification, the cancer cell profile (epithelial Luminal, epithelial Basal, epithelial HER2) with the highest proportion was used to reclassify the sample. The TCGA annotations were used to determine the PAM50 classification of the TCGA tumors.

For the cell line XDec-SM classification, the BRCA cell lines from the Cancer Cell Line Encyclopedia were self-self-clustered with the epithelial profiles obtained from the deconvolution of TCGA over the PAM50 genes. Four distinct groups of cell lines were identified that matched the *in vivo* epithelial profiles. The PAM50 classifications were obtained from previous literature (Jiang et al., 2016).

##### Tamoxifen Survival Analysis

The recount package was used to determine the treatment of TCGA models. 72 tumors were identified that were treated with tamoxifen and classified as Luminal by PAM50. We then compared the tumors that were also classified as Luminal by XDec-SM vs those that were classified as HER2 by XDec-SM using the survival and survminer packages in R.

##### PDX Datasets

BCM cohort: A set of 50 TNBC PDX models were obtained from the BCM PDX collection, which are available on the PDX portal (https://pdxportal.research.bcm.edu/). These models were treated with four cycles of human equivalent docetaxel (20mg/kg, IP), carboplatin (50mg/kg, IP) and the control models were untreated for four weeks, and response was measured quantitatively as the change in tumor volume from baseline. Deep RNA-seq was also obtained for all models in this collection (∼200M reads/sample). The murine vs human reads were separated by Xenome and the deconvolution was done on the human reads. Of the 50 models, 44 models that didn’t model the same patient had a combined epithelial proportion of > 0.7 and an epithelial Basal + epithelial HER2 + epithelial Luminal proportion of > 0.1 and were used for the response predictions.

RMGCRC cohort: The RNA sequencing and response to therapy for 30 PDX models from the RMGCRC were obtained from GEO (GSE142767). The response for these models was classified according to the RECIST criteria. Of these models, 22 had a combined epithelial proportion of > 0.7 and an epithelial Basal + epithelial HER2 + epithelial Luminal proportion of > 0.1 and were used for the response predictions.

##### Response To Chemotherapy

PDX: A GLM model was built on the BCM dataset for carboplatin and docetaxel using all models from the epithelial Basal, epithelial Luminal, and epithelial HER2 proportions. These GLMs were then applied to the RMGCRC dataset for drug treatments in the same class (cisplatin and carboplatin, paclitaxel, and docetaxel) and an ROC was calculated for qualitative response.

## SUPPLEMENTAL INFORMATION

**Figure S1. XDec-SM Deconvolution of RNA-seq profiles of TCGA breast tumor samples utilizing scRNA-seq references, related to Figure 1, Figure 2, Figure 3, STAR Methods.**

(A) Heatmap representing the transformed gene expression counts of the 274 informative genes across the pseudo bulk reference expression profiles generated from the scRNA-seq gene expression profiles.

(B) Heatmap representing the correlation between the XDec-SM estimated expression profiles (n = 9) and the pseudo bulk reference expression profiles. Red boxes are placed over the highest correlation. XDec-SM estimates five epithelial profiles, two stromal profiles, one T cell profile, and one macrophage profile.

(C) Heatmap representing the per-sample proportion of the nine constituent cell types in the TCGA BRCA dataset. Top color bar represents the PAM50 expression subtypes. Only samples with identified subtypes are included in the heatmap.

(D) Boxplot representing the per-sample proportion of the nine constituent cell types in the TCGA BRCA dataset separated by the PAM50 classification subtypes.

**Figure S2. Cell type specific gene expression, related to STAR Methods.**

The cell type specific gene expression of marker genes across different breast cancer subtypes. ESR1 and FOXA1 are most highly expressed in the epithelial compartment. ADIPOQ and FABP4 are most highly expressed in the stromal adipocyte compartment. FN1 and COL1A1 are most highly expressed in the stromal CAF compartment. CD3G and CD8A are most highly expressed in the immune compartment.

**Figure S3. Distribution of scRNA-seq profiles for individual tumors, related to Figure 3.**

(A-H) Single cell RNA-seq and bulk RNA seq for the same tumor plotted on the cancer cell state map.

**Figure S4. Survival curves for patients treated with Tamoxifen, related to Figure 6.**

(A) Survival curve showing no significant survival difference between patients classified as Luminal by PAM50 and HER2 negative (blue) and Luminal by PAM50 and HER2 positive (pink).

(B) Survival curve showing no significant survival difference between patients classified as Luminal by PAM50 and with high ERBB2 pathway activation (blue) and Luminal by PAM50 and without high ERBB2 pathway activation (pink).

