## Supplementary Material for "XDec Simplex Map of Breast Cancer Cell States Enables Precise Modeling and Targeting of Breast Cancer"

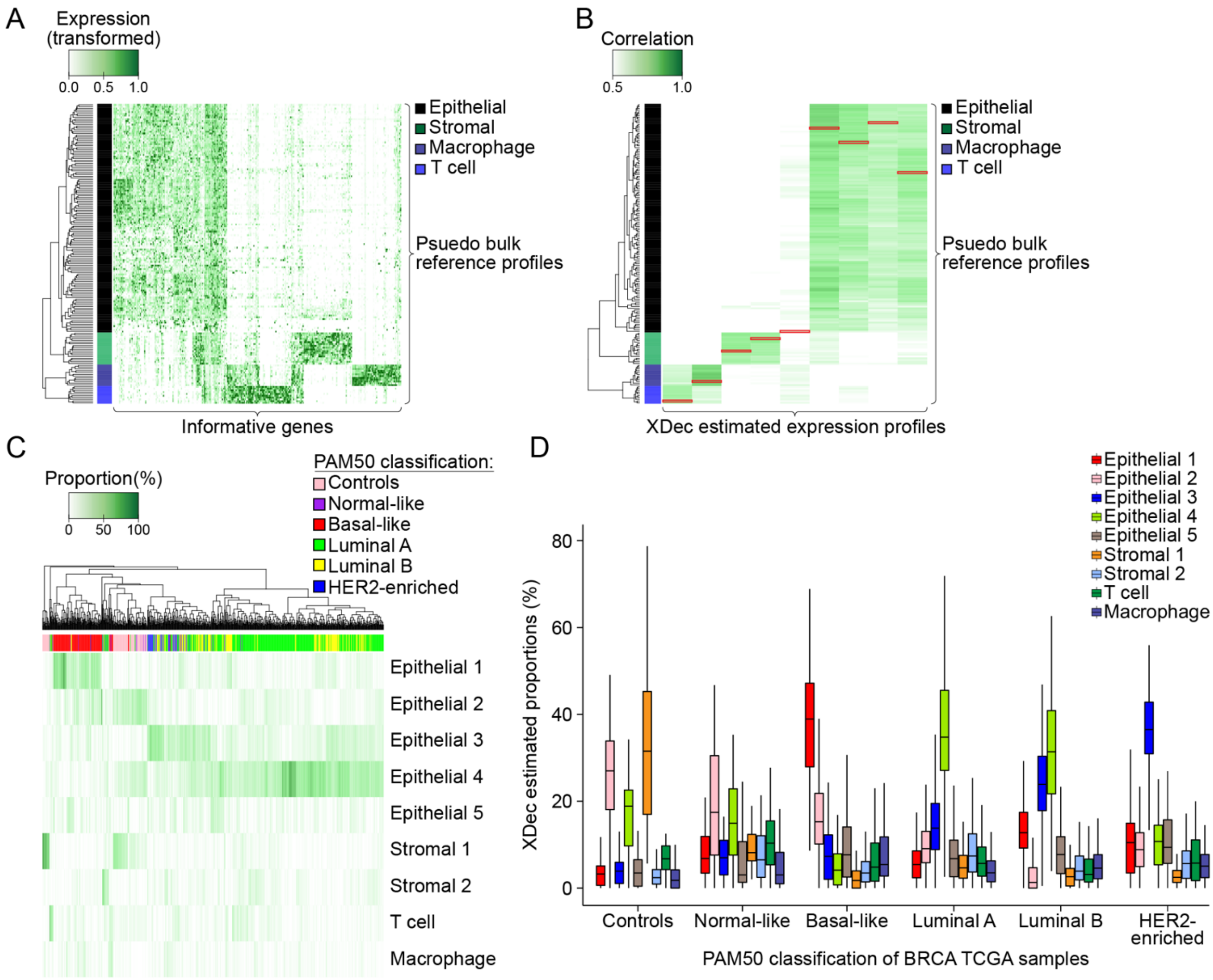


**Figure S1. XDec-SM Deconvolution of RNA-seq profiles of TCGA breast tumor samples utilizing scRNA-seq references**

(A) Heatmap representing the transformed gene expression counts of the 274 informative genes across the pseudo bulk reference expression profiles generated from the scRNA-seq gene expression profiles.

(B) Heatmap representing the correlation between the XDec-SM estimated expression profiles (n = 9) and the pseudo bulk reference expression profiles. Red boxes are placed over the highest correlation. XDec-SM estimates five epithelial profiles, two stromal profiles, one T cell profile, and one macrophage profile.

(C) Heatmap representing the per-sample proportion of the nine constituent cell types in the TCGA BRCA dataset. Top color bar represents the PAM50 expression subtypes. Only samples with identified subtypes are included in the heatmap.

(D) Boxplot representing the per-sample proportion of the nine constituent cell types in the TCGA BRCA dataset separated by the PAM50 classification subtypes.

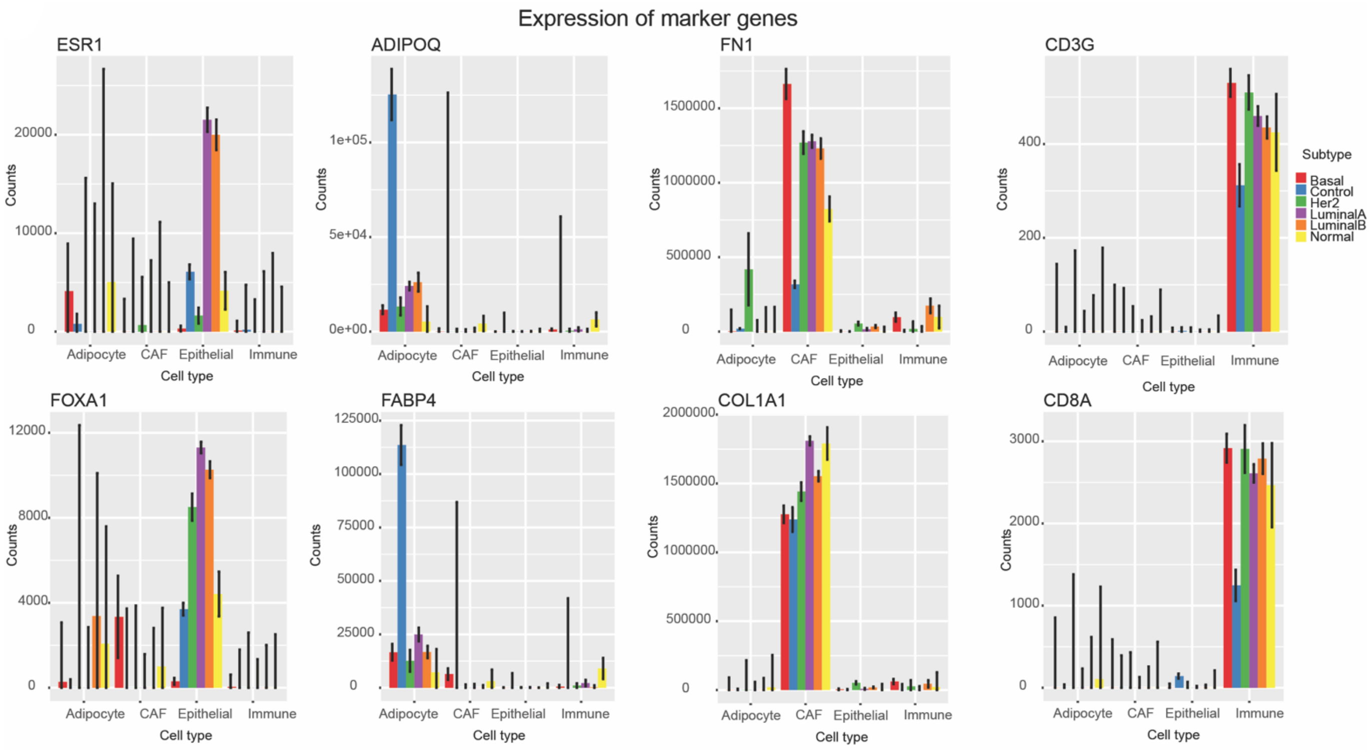


**Figure S2. Cell type specific gene expression**

(A) The cell type specific gene expression of marker genes across different breastcancer subtypes. ESR1 and FOXA1 are most highly expressed in the epithelial compartment. ADIPOQ and FABP4 are most highly expressed in the stromal adipocyte compartment. FN1 and COL1A1 are most highly expressed in the stromal CAF compartment. CD3G and CD8A are most highly expressed in the immune compartment.

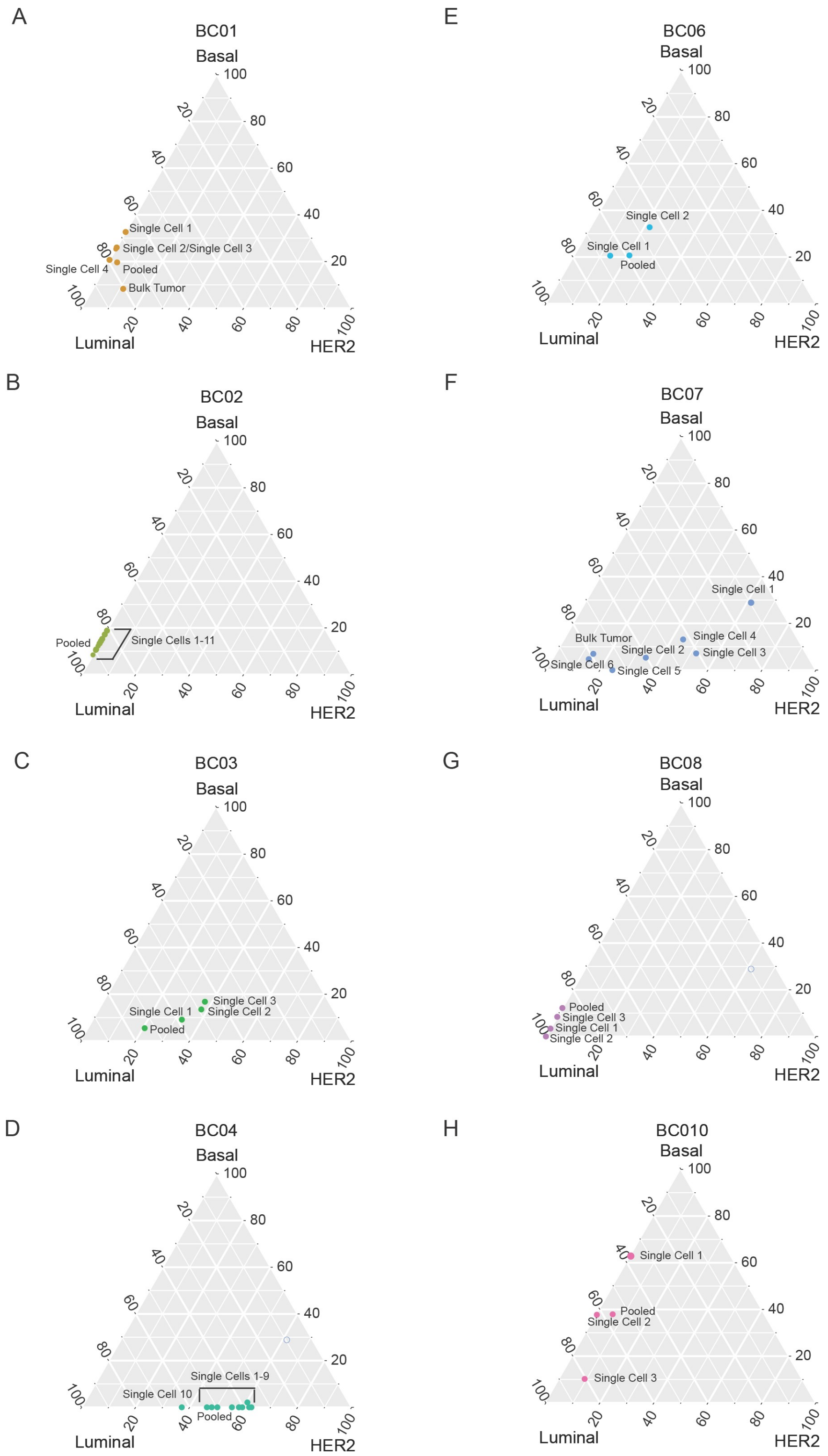


**Figure S3. Distribution of scRNA-seq profiles for individual tumor**

(A-H) Single cell RNA-seq and bulk RNA seq for the same tumor plotted on the cancer cell state map.

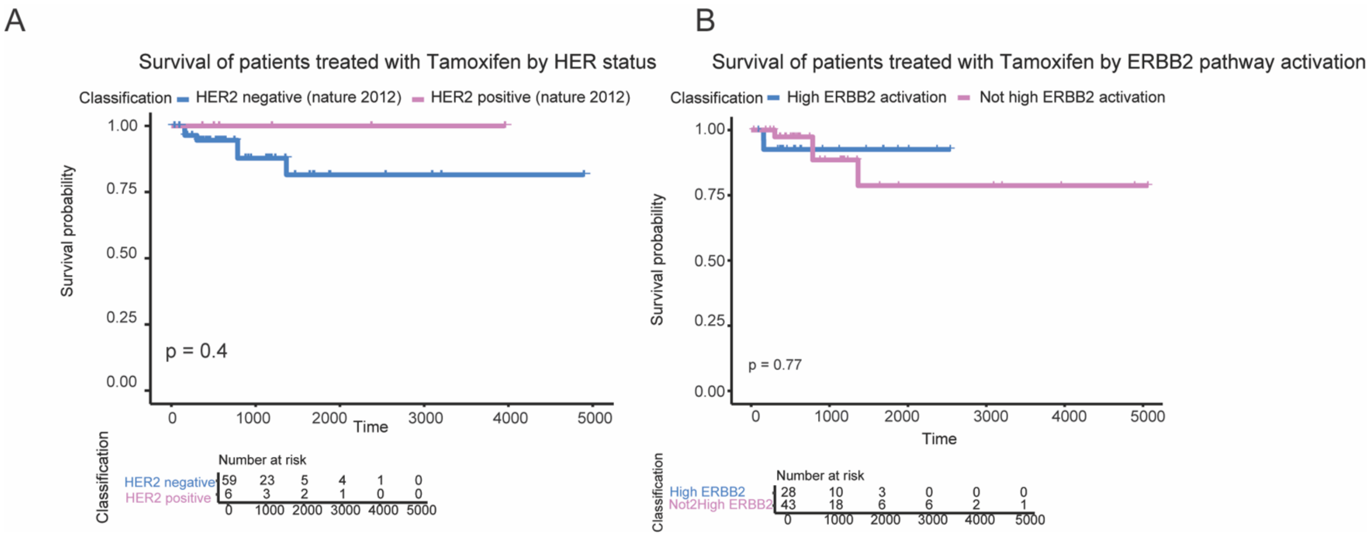


**Figure S4. Survival curves for patients treated with Tamoxifen**
(A left) Survival curve showing no significant survival difference between patients classified as Luminal by PAM50 and HER2 negative (blue) and Luminal by PAM50 and HER2 positive (pink). (A right ) Survival curve showing no significant survival difference between patients classified as Luminal by PAM50 and with high ERBB2 pathway activation (blue) and Luminal by PAM50 and without high ERBB2 pathway activation (pink).
